# Natural thermal stress-hardening of corals through cold temperature pulses in the Thai Andaman Sea

**DOI:** 10.1101/2023.06.12.544549

**Authors:** Marlene Wall, Talisa Doering, Nina Pohl, Lalita Putchim, Tipwimon Ratanawongwan, Anna Roik

## Abstract

Thermal variability can render corals stress resistant through a phenomenon coined as “stress-hardening induced by environmental priming”. Fluctuations that involve high temperature peaks have been commonly investigated, however, the effects of a stress-hardening stimulus generated by cold-water pulses has rarely been studied. Offshore island reefs in the Andaman Sea offer an ideal natural setting to study these effects, as cooling water of internal waves induce strong variability with peak intensity in January to June and absence in August to November. While western island shores are exposed to this stimulus, eastern shores remain sheltered. This study examined (1) whether corals from exposed reefs were more heat stress resistant compared to stimulus-sheltered conspecifics and (2) whether this trait can last in the absence of the stimulus. We quantified the thermal stress resistance in two ecologically important coral species, *Pocillopora* sp. and *Porites* sp., from the two island shores, during the two seasons. Coral bleaching intensity and photosynthetic efficiency of algal symbionts were measured as response variables after a short-term heat stress assay (24-48 h, 34 °C) to assess thermal stress resistance. Stress responses of all stimulus-exposed corals were either undetectable (during the season of stimulus presence) or very weak (during stimulus absence), while corals from the stimulus-sheltered shore responded strongly to heat stress irrespective of the season. Hence, thermal resistance was overall greater in corals originating from the stimulus-exposed shore, but it was slightly diminished during the season of stimulus absence, emphasizing the relevance of stimulus recurrence in maintaining the resistance trait. We exemplify that the stimulus of fluctuating low temperature pulses successfully induced stress-hardening in corals. This suggests that priming stimuli do not necessarily need to transgress certain upper thermal thresholds, but can also touch on lower thresholds to be effective. Even more, we argue that cooling pulses might represent a safer stress-hardening regime, since warming-stress accumulation can be avoided. More research is required to obtain a better understanding of environmental priming, but current findings should encourage the development of artificial stress-hardening approaches to enhance coral resistance in reef restoration efforts.

## Introduction

Reef-building corals live near their thermal limits, so that the growing thermal stress caused by ocean warming poses the most pressing threat to the existence of many coral species and the tropical reef ecosystems (Hoegh-Guldberg 1999). Thermal anomalies that culminate during long-lasting heat-waves (Sully et al. 2019) impose intense stress on corals, which triggers “coral bleaching”, i.e., the loss of symbiotic algae from the coral holobionts that is apparent through the paling of the coral tissues (Glynn 1991). Bleached corals are exposed to starvation, as the nutrient exchange between the host and dinoflagellate symbionts is disrupted (Brown 1997; Rädecker et al. 2021). As a result, coral bleaching events have already been fatal to vast portions of coral populations worldwide, resulting in significant losses of the coral reef ecosystem (Hughes et al. 2018).

Tropical corals have adapted to mostly stable thermal conditions of the tropical waters which feature only little seasonal change (Kleypas et al. 1999). Consequently, they do not cope well with even slight increases in temperature. However, observations of coral reef habitats that feature comparably high thermal (and other) fluctuations provide a glimpse into the remarkable plasticity of certain individuals or populations. Such studies have demonstrated that corals, pre-exposed to challenging conditions, can feature a higher thermal resistance, especially in terms of their thermal threshold and/or show a higher ability to recover after (thermal) stress compared to their counterparts living in more stable environments nearby. Subsequently, this phenomenon has been coined “environmental priming” or “environmental memory” (Brown et al. 2002; Rivest et al. 2017; Hackerott et al. 2021; Martell 2023). Most of these observations originate from reefs that experience strong environmental variability, e.g., inshore reef habitats (Kenkel and Matz 2016), reef flats and tidal pools that are temporarily exposed to extreme conditions (Oliver and Palumbi 2011a; Schoepf et al. 2015), sites exposed to internal waves or other upwelling events (Doering et al. 2021; Buerger et al. 2015), or reef-adjacent habitats, such as lagoons and mangroves (Camp et al. 2016, 2017). Also, temporal stress events, such as moderate heat waves (Bellantuono et al. 2012b; Ainsworth et al. 2016; Fox et al. 2021) or consecutive bleaching events (Maynard et al. 2008; Guest et al. 2012; Penin et al. 2013), have been observed to be associated with greater stress resistance and/or a faster recovery of corals following such events, where mechanisms underlying adaptation and/or acclimation can be suspected.

The phenomenon of stress resistance gain is not unique to corals. Stress-hardening through environmental priming relies on the phenotypic and physiological plasticity of an organism (Hilker et al. 2016) and has long been observed and studied across many taxa of the tree of life, most prominently in plants (Nicotra et al. 2010; Tanou et al. 2012; Li et al. 2014). Past environmental challenges can “prime” organisms to respond to future stressors more efficiently and/or rapidly. In contrast to the mechanisms of adaptation, which arise from genetic variation and selection dynamics over several generations, the effect of environmental priming can occur within one generation and in the same individual (Whitman and Agrawal 2009; Foo and Byrne 2016). For long-lived coral species with relatively long generation times and, hence, naturally slower adaptive processes, rapid acclimation through plasticity can become life-saving (Palumbi et al. 2014). As such, environmental priming could fundamentally increase the odds for corals to successfully resist rapid ocean warming.

Today, reefs that exhibit high thermal variability have become attractive sites to obtain stress tolerant corals to study the mechanisms underpinning plasticity and thermal resistance (Oliver and Palumbi 2011b; Ziegler et al. 2017; Hackerott et al. 2021; Majerova et al. 2021). Furthermore, study of these stress resistant phenotypes can assist the development of interventions to enhance coral stress tolerance (Doering et al. 2021; Epstein et al. 2019; Howells et al. 2021). Most recently, *ex situ* experiments aiming to develop stress-hardening procedures for the laboratory have gained traction. Such studies have shown that artificial preconditioning treatments applying fluctuating temperatures in artificial aquarium environments, can improve thermal stress resistance of corals (Hawkins and Warner 2017; Majerova et al. 2021; Alexander et al. 2022; DeMerlis et al. 2022). However, findings are still equivocal, as several other studies did not report any stress-hardening effects (Putnam and Edmunds 2011; Klepac and Barshis 2020; Alexander et al. 2022; Schoepf et al. 2022). Obtaining an understanding of the greater detail of these phenomena is of urgent relevance. Hence, important questions concerning the “dosage” of a priming stimulus, including the exposure duration and the regime of the priming conditions which is required to achieve the desired effect, remain to be answered (Brown et al. 2023; Martell 2023).

To shed light on some of the questions, this study took advantage of the environmentally and seasonally dynamic reef sites located in the Andaman Sea in Thailand, where island shores are seasonally exposed to a stimulus of thermal variability induced by large amplitude internal waves (Osborne and Burch 1980). Internal waves are oceanographic features that are ubiquitous in the world’s oceans and, contrary to surface waves, travel deep along strong density gradients (Jackson et al. 2012). Particularly in the Thai Andaman Sea, western reef sites are exposed to the internal waves and experience the strongest impacts. Internal waves induce variations of temperature (i.e., negative anomalies with minima at 26 to 24 °C), pH, salinity and other environmental variables during their peak season of wave intensity that is from January to June. They can impose tremendous stress on corals, but can provide protective cooling during natural marine heat waves (Wall et al. 2015; Wyatt et al. 2019). In contrast, reefs on the east shores of the islands are sheltered from these impacts, fostering stable environmental conditions (Schmidt et al. 2012; Wall et al. 2012). During the second half of the year, impacts of internal waves are typically minor or absent on the western shores, which allows the study of coral thermal stress resistance in presence and absence of the priming stimulus. While a majority of studies have focused on coral stress-hardening by exposing them to high temperature regimes under constant or variable conditions (Middlebrook et al. 2008; Bellantuono et al. 2012b; Putnam and Gates 2015; Hackerott et al. 2021; Martell 2023), internal wave sites provide the opportunity to study the effect of high variability conditions that do not involve high but rather low thermal pulses.

A previous study has established that internal wave impact, in the Andaman Sea region, was linked to higher coral thermal stress resistance, specifically in *Porites* sp. corals studied during the season of high internal wave intensity (Buerger et al. 2015). We followed up on these previous findings and investigated whether this stress-hardening effect reported for *Porites* sp. is a species-specific phenomenon or can be also found in other coral species. Second, we investigated whether a greater thermal resistance of corals originating from the stimulus-exposed reefs is a persistent or a transient trait, which occurs only during the season when environmental variability is at its peak. To address these questions, we employed high-throughput short-term heat stress assays (duration: 24 - 48h, peak of 34 °C) (Doering et al. 2021; Evensen et al. 2021) to assess the thermal stress responses in two coral species, *Pocillopora* sp. and *Porites* sp., from a stimulus-exposed and a sheltered site during the two seasons.

## Materials and Methods

### Study sites and coral collection

Study sites were located at Racha Island in the Andaman Sea off the coast of Thailand, both at 15 m water depth (Figure 1 A-B). A reef on the western shore was chosen (7.595530°N, 98.354320°E, Figure 1 B) where internal wave forcing induced environmental variability through frequent upwelling of deep, cool, and nutrient rich water onto the shelf (Wall et al. 2012; Schmidt et al. 2016). A reef on the eastern shore, sheltered from the internal wave stimulus, was chosen to represent a low variability reef (7.598910°N, 98.373100°E, Figure 1 B). Temperature fluctuations were monitored *in situ* as a proxy for internal wave impact and environmental variability. Temperature loggers (HOBO Pendant Temperature/Light 8K Data Logger, Onset, USA) were deployed at the study sites one month before heat stress assays were performed. In each study site, visually healthy coral colonies of *Pocillopora* sp. and *Porites* sp. were permanently tagged to assess their thermal resistance levels during the two seasons (n=8 to n=18, Figure 1 C, Table S1). These two coral species are cosmopolitan reef-builders in Thailand and within the entire Indo-Pacific region (Brown and Phongsuwan 2012; Schmidt et al. 2012; Jain et al. 2023). Coral fragments were collected at the end of April 2018, an episode of strongest internal wave impact, and at the end of October (*Porites* sp.) and November (*Pocillopora* sp.), during the absence of internal wave stimulus. Two fragments (*Porites* sp.: ø ∼ 6 cm; *Pocillopora* sp.: length ∼ 5 cm) per colony were collected using a chisel and a hammer (Table S1).

**Figure 1.**
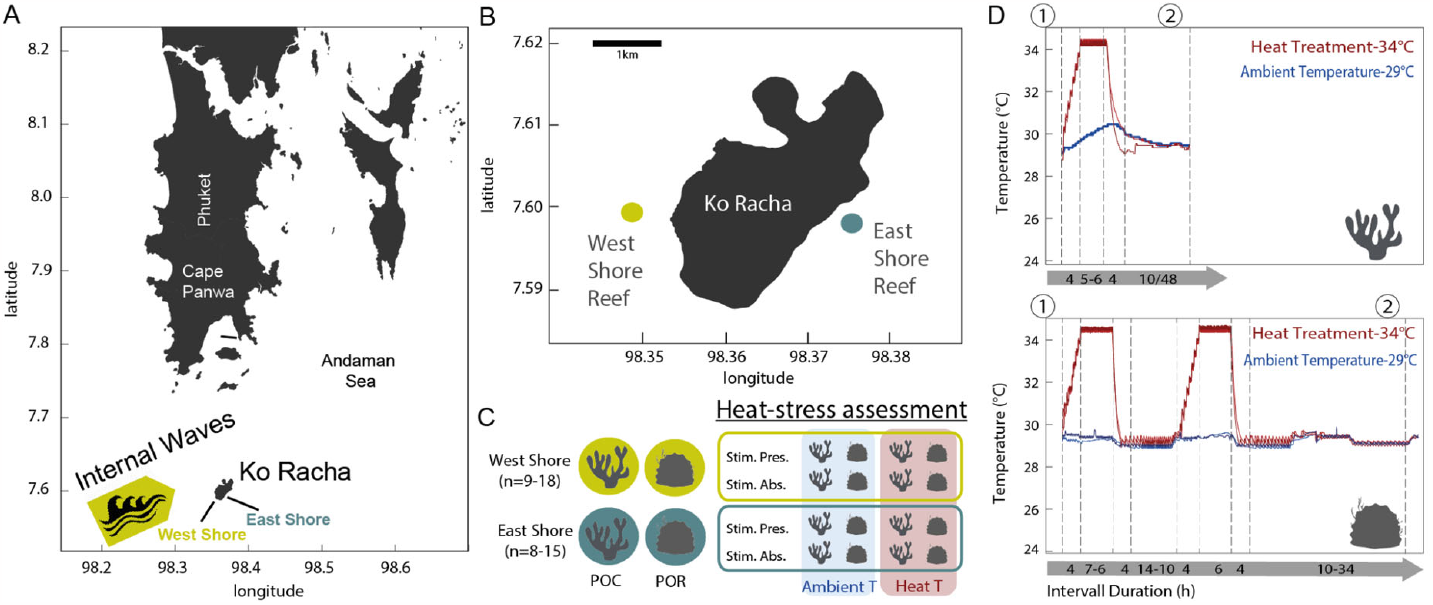
Study sites, experimental design, and temperature profiles of short-term heat stress assays. (A) The study area was located at Ko Racha in the Andaman Sea off the coast of Thailand, a region that is exposed to large amplitude internal waves (light green arrow). These deep waves cause high thermal (and environmental) variability at western reef sites (light green), while the eastern shores (dark green) remain sheltered. (B) Two reef sites were chosen, on each shore side. (C) Fragments of colonies (n = replicate numbers) of *Pocillopora* sp. (POC) and *Porites* spp. (POR) were collected for heat tolerance assessment during two seasons - the season of stimulus presence and peak of internal wave frequency and intensity (Stim. Pres.) and peak of internal wave frequency and intensity in April and the season of stimulus and internal wave absence (Stim. Abs.) in October/November. Fragments were subjected to a short-term heat stress assay exposing them to a heat pulse treatment of 34 °C (Heat T). A control group was maintained at ambient temperature of 29 °C (Ambient T). (D) Temperature profiles were customized for each coral species, accounting for their taxon-specific temperature sensitivity. Pocilloporid fragments were challenged with a single heat stress pulse over one day (upper panel, branching coral icon), while *Porites* sp. fragments required two heat stress pulses over two days to show a heat stress response (lower panel massive coral icon). Measurement timepoints at the start (1) and the end (2) of each experiment are indicated.

### Short-term heat stress assays

Collected fragments were instantly transported to the Phuket Marine Biological Center (Phuket, Thailand) where they were maintained in two 500 L flow-through tanks with a flow rate of 2.8 ± 1.31 L/min until the start of each heat stress assay. Another 500 L source tank constantly supplied both flow-through tanks with 5 μm-filtered seawater from the reef adjacent to the research center. Its temperature was held at constant 29.43 ± 0.32 °C using a temperature-controlling device including a chiller and a heater (Titanium Heater 100 W, Schego, Germany; Temperature Switch TS 125, HTRONIC, Germany; Aqua Medic Titan 1500 Chiller, Germany). LED lights (135 W, Hydra Fiftytwo HD LED, Aqua Illumination, USA) mimicked the average light conditions of the sampling sites (Text S1).

For each heat stress assay (Figure 1 D), two 40 L experimental tanks were set up inside each of the 500 L flow-through tanks that were used as temperature-controlling water baths (Table S2). The seawater of all four experimental tanks was supplied by daily, manual 50% water changes from the source-tank. Each experimental tank was equipped with a temperature-controlling device, one heater, air supply, a small current pump and a temperature logger (Temperature Switch TS 125, HTRONIC, Germany; Titanium Heater 100 W, Schego, Germany; Koralia nano 900 L/h, Hydor, Italy; HOBO Pendant Temperature/Light 8K Data Logger, Onset, USA). Two coral fragments per coral colony were randomly distributed among the four tanks “34°C” (*N* = 2) and “29°C” (*N* = 2), resulting in one fragment per colony per treatment. The 34°C-treatment was established over the course of one day by ramping temperatures from 29°C to 34°C for 4 h, holding at 34°C for 5 h or 6 h (*Pocillopora* sp.) or for 6 h or 7 h (*Porites* sp.), and decreasing temperatures to 29 °C within 4 h. After the heat exposure, corals were maintained at ambient temperatures for 10 h until the next day. While *Pocillopora* sp. fragments were subjected to the short-term heat exposure once, resulting in a 24 h experiment, *Porites* sp. corals were exposed to the treatment over two consecutive days resulting in a duration of 72 h (Figure 1 D).

### Coral stress response variables

We measured two variables that assessed the stress response of each fragment before and after each heat stress assay (timepoints (1) and (2) in Figure 1 D). Tissue coloration, a proxy for microalgal symbiont cell density in coral tissues and therefore an indicator of holobiont health and coral bleaching severity, was assessed using a “bleaching score”. The coloration of each individual fragment was visually categorized on the scale from 1 (bleached, pale tissues) to 6 (healthy, dark tissues) using a coral bleaching chart (Siebeck et al. 2006). A minimum and maximum score was recorded per fragment and averaged. Photosynthetic efficiency of microalgal symbionts was determined by measuring effective quantum efficiency (yield Φ PSII = (Fm’ – F) / Fm’ = ΔF / Fm’, Genty et al. 1989) of electron transport using a pulse amplitude-modulated fluorometer (Diving-PAM, Walz, Germany).

### Statistical analyses

Δ-values of each stress response variable (end – start of each experimental part) were calculated to represent the change or the variable over time. Based on these Δ-values, effect sizes were estimated using *dabestR* v0.2.3 6 (Ho et al. 2019). Effects of the high temperature treatment (“34 °C” vs. “29 °C”) were compared between the sites of origin (“West | High variability site” and “East | Low variability site”) and between the seasons (“Season of stimulus presence” and “Season of stimulus absence”). Statistical significance was tested in *R* (R Core Team 2013) using linear mixed effect models (*nlme* v4 3.1-148 and *lme4* v1.1-23 package). Where applicable, coral colony genotype was used as a random factor.

## Results

### Environmental variability of the study sites

Temperature was recorded as a proxy for internal wave forcing and provided a measure of environmental variability on the study sites. The temperature profiles revealed that the intensities of internal waves were seasonal (Figure 2). Strong and mainly negative temperature anomalies occurred during March to April, which provided the strongest stimulus for environmental priming with a diurnal amplitude ranging between 0.7 - 5.4 °C (average amplitude of 3.0 °C ± 1.0 SD) and minimal temperature values as low as 24 °C (Figure 2 A, C). The impact of internal waves dwindled in September to November, when the anomalies decreased (Figure 2 B, C). Importantly, temperature anomalies driving the environmental variability were more frequent and intense on the exposed west shore of Ko Racha compared to the sheltered east shore (Figure 2 C). During the season when the stimulus of internal waves was present, the differences were largest between both island sides, east and west (Figure 2 A). Once the stimulus faded during the second half of the year, conditions on both island sides became more similar (Figure 2 B).

**Figure 2.**
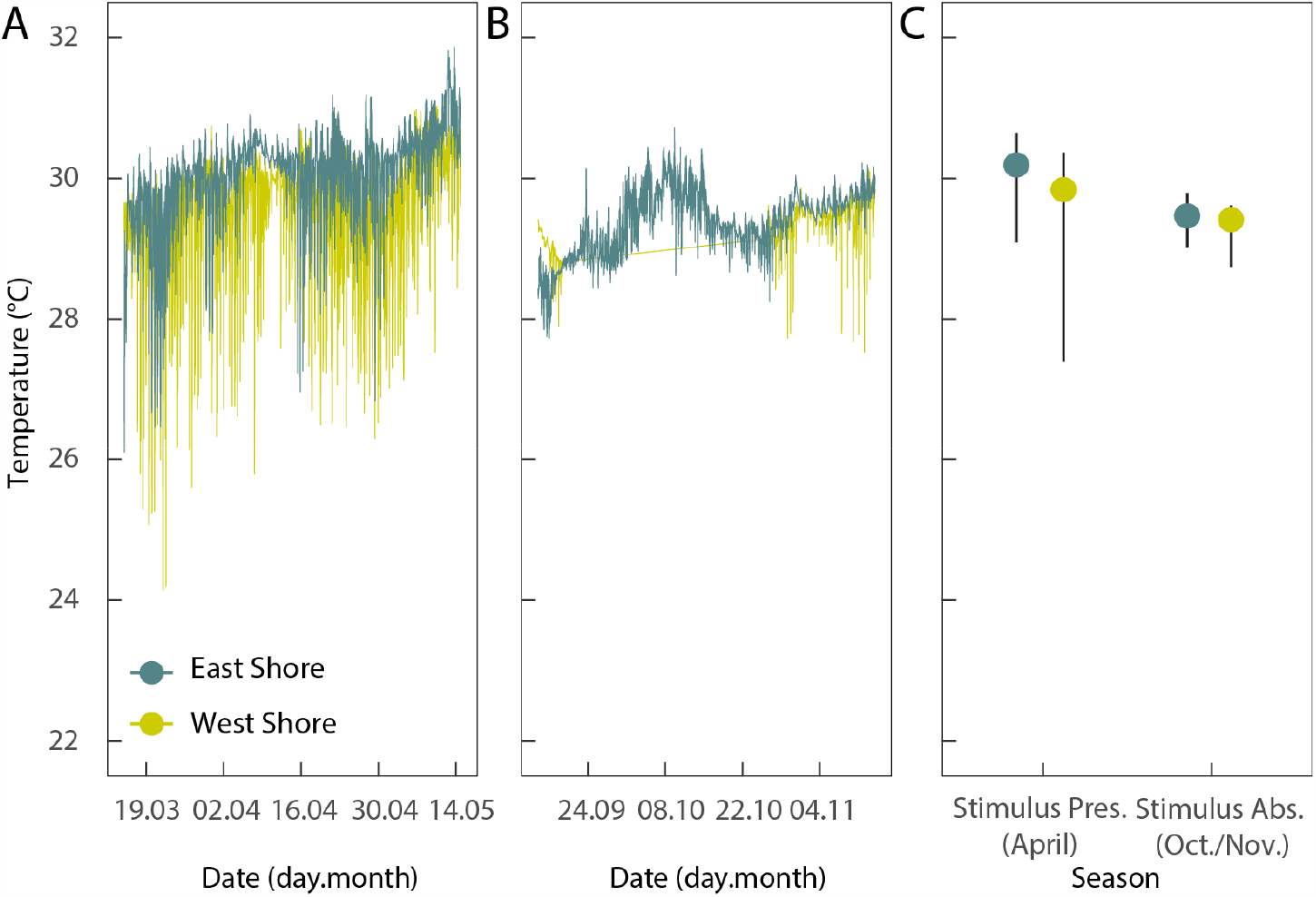
Seasonal difference of environmental variability at the study sites. Temperature records served as a proxy for environmental variability in the study sites. This variability was generated by large amplitude internal waves, a potential stimulus for coral stress-hardening. (A) A time series depicts that internal waves on the western shores (light green) induced strong temperature anomalies during March to April. The eastern shore (dark green) remained mostly sheltered from this stimulus. (B) Fairly constant temperatures, with almost similar dynamics on both island shores, were characteristic for the second part of the year (October to November), the season of stimulus absence. During this time, the western and eastern shores featured more similar conditions. (C) The median temperatures are indicated by circles and the average positive and negative diurnal anomalies from the median are displayed as whiskers for both island shores and seasons.

### Stress responses to short-term heat stress assays

Overall, the bleaching score and photosynthetic efficiency data indicated that stress levels after the short-term heat stress assay were highest in corals from the eastern, stimulus-sheltered site, as reflected in the lighter color tones in the heatmap, where effect sizes are visualized (Figure 3). This becomes clear, as the largest and significant declines in the two variables, tissue coloration and photosynthesis, were recorded in east shore corals irrespective of the season. Corals from the west shore did not show any significant signs of stress when tested during stimulus presence (Figure 3A), but significant declines of the two variables were noted, when corals were tested during stimulus absence (Figure 3B). Here effect sizes of the heat stress treatment were fairly small (< 0.9) for tissue coloration, indicating a mild stress response. In comparison, the large stress responses measured for the east shore corals were in the effect size range of 1.05 - 1.97 (for the tissue coloration). Interestingly, the decline in photosynthesis of the west-shore corals during stimulus absence was comparable to the decline of photosynthesis in corals from the east shore.

**Figure 3.**
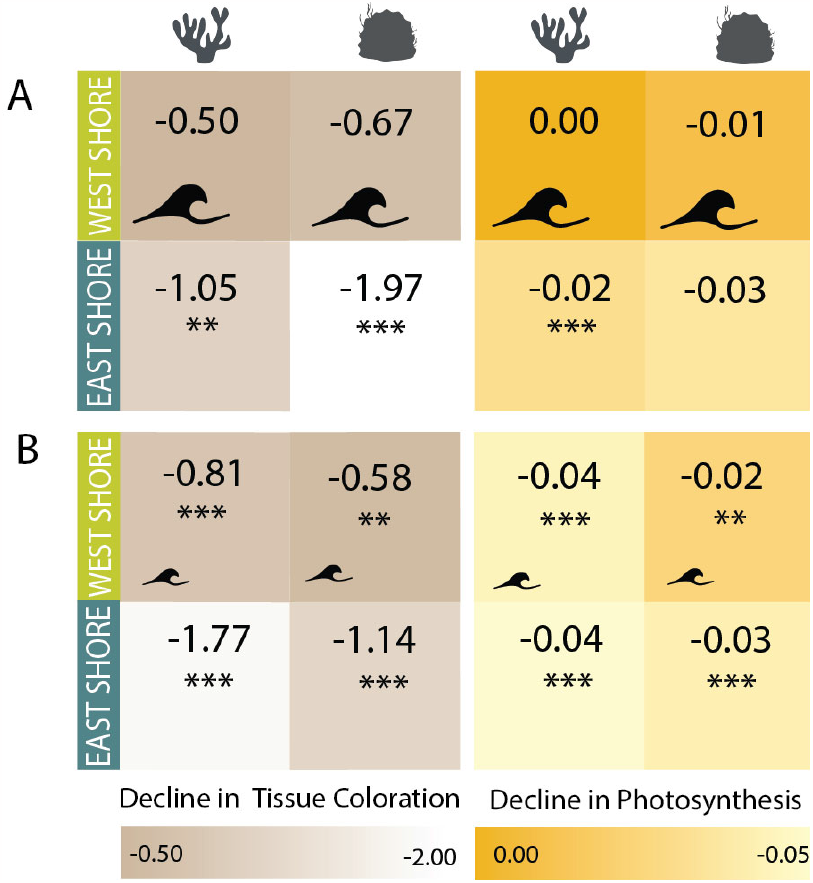
Summary of thermal stress levels of corals under experimental heat exposure compared between their sites of origin and seasons. Stress levels of corals assessed in short-term heat stress assays during the (A) season of stimulus presence (April) and the (B) season of stimulus absence (November) are visualized. Lighter tones indicate higher stress levels. Colors represent the effect sizes determined using Cohen’s d metric as the mean differences of measurements between the heat treatment and ambient control group of the heat stress assays. Negative values and lighter color tones indicate decreases of the bleaching score (brown tones) and the decline in photosynthetic efficiency (yellow tones) as a result of heat stress. Significant effects are marked as *p* <0.001***, < 0.01**, <0.05* as obtained from the post hoc tests of generalized linear mixed models. Dark green= eastern sheltered shore; light green = western exposed shore; the size of the wave icon indicates the magnitude of internal wave impact on the reef as a stress-hardening stimulus.

#### Bleaching responses

Across the seasons the bleaching score of corals from the eastern sheltered reef strongly declined under the acute heat exposure during the heat stress assay, as indicated by significant loss of tissue coloration (29 °C group vs. 34 °C group, *p* < 0.001, Figure 4 A-D, Tables S3-4 and S6). Negative effect sizes were largest in these east-shore corals, i.e., -1 to -2, which was mostly 2 to 4-fold larger, compared to those of corals from the western reef (i.e., effect sizes of 0 to -0.8). In contrast, the stress responses of westshore corals differed between the seasons. Overall, their bleaching score did not decline in response to heat stress, when assessment was conducted during the season of stimulus presence (Figure 4 A, C, Table S6). However, during the second half of the year (i.e., stimulus absence season), tissue coloration of corals from the western shore slightly declined under experimental heat exposure with rather small, but measurable, differences between the heat and ambient temperature control group (Figure 4 B, D, Table S6). A small but significant decrease of the bleaching score was recorded (*p* < 0.001, Figure 3 B, D). Further, a small-scale decline in the bleaching score was recorded for *Porites* sp. assessed during both seasons with effect sizes were around ∼0.6 (*n*.*s*. under stimulus presence and *p* < 0.01 under stimulus absence, Figure 4 C-D, Table S4, S6).

**Figure 4.**
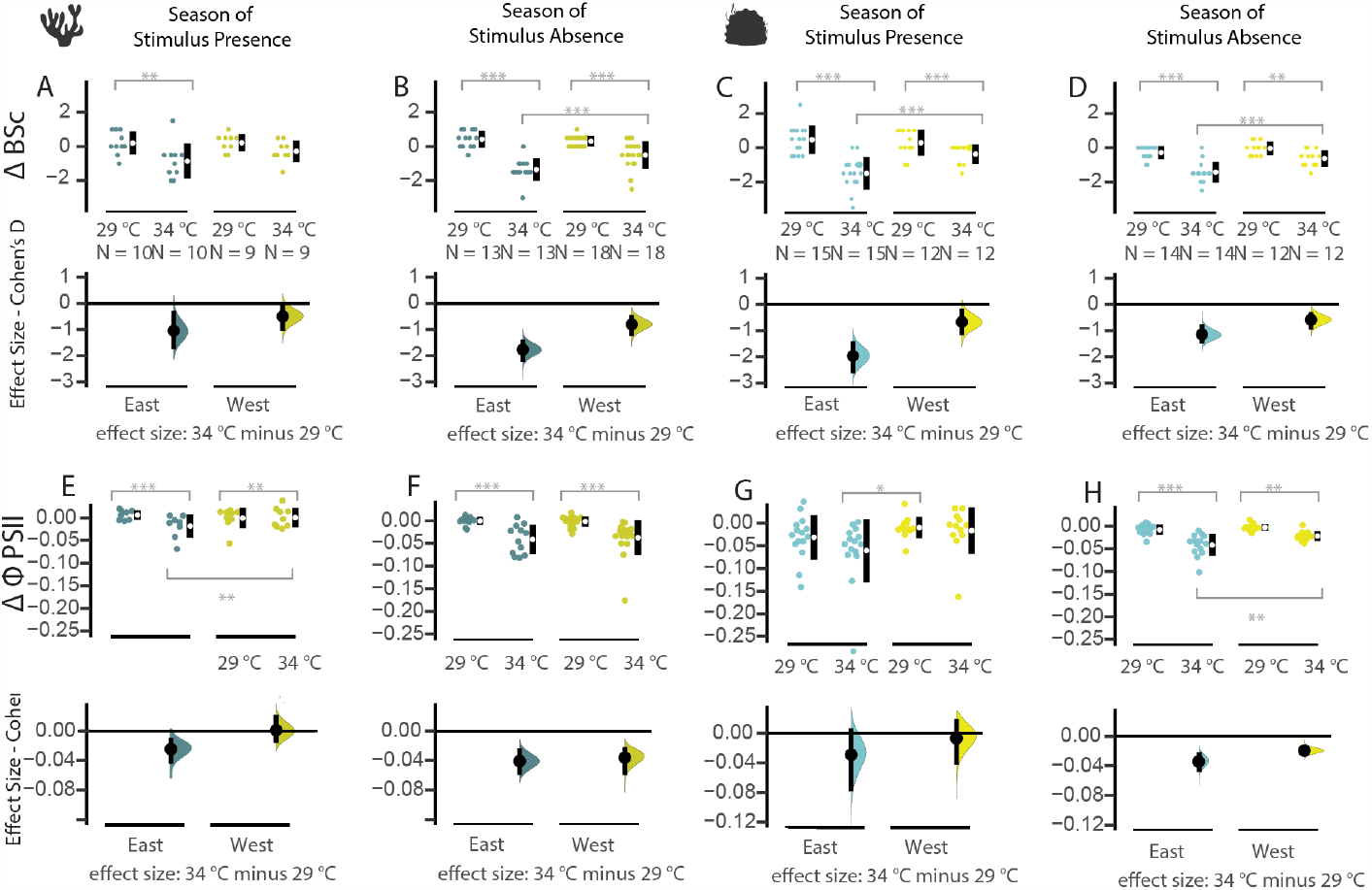
Symbiont loss and photosynthetic efficiency change of corals under experimental heat exposure compared between their sites of origin and seasons. Estimation plots compare the effects of acute experimental heat exposure (“29°C” vs. “34°C”) on (A - D) the loss of coral tissue coloration (bleaching score = “BSc”) and (E - H) the photosynthetic efficiency of coral microalgal symbionts (quantum efficiency of electron transport = Δ-Φ PSII). Data is shown for the corals *Pocillopora* sp. (“branching” coral icon) and *Porites* sp. (“massive” coral icon) during the two seasons of stress-hardening stimulus presence and absence. The data is presented as a Δ-value of the measured variables (i.e., the decline of values between the start and end of stress exposure). Negative values indicate the loss of pigments and microalgal symbiont cells from the coral tissues or the decline of photosynthetic efficiency as a result of heat stress. Swarm plots show raw data points (first and third row) and Cumming Estimation plots (second and fourth row) depict the effect sizes as the mean differences between the experimental groups using Cohen’s d and a 95% confidence interval. Significant differences between the stress responses within the groups are indicated by connecting lines (p <0.001***, < 0.01**, <0.05* obtained from the post hoc tests of generalized linear mixed models). Vertical error bars = 95% CI; *N* = individuals per treatment group.

#### Declines in photosynthetic efficiency

Across the two seasons, photosynthetic efficiency decreased significantly after the heat stress exposure in almost all coral fragments after heat exposure (29 °C group vs. 34 °C group, *p* < 0.001, Figure 4 E-H, Tables S3, S5, S8), except for *Porites* sp. assessed during the season of stimulus presence (*n*.*s*., Figure 4 G, Table S8). Despite a significant response of almost all corals, the declines of photosynthetic efficiency measured by effect size were larger in corals from the eastern reef, i.e., -0.02 to -0.03, compared to corals from the western reef, i.e., mostly 0 or -0.01 (e.g., *p* < 0.01, Figure E and H).

## Discussion

This study investigated the dynamics of coral thermal resistance in relation to a seasonally changing priming stimulus of internal waves in the Thai Andaman Sea. Our data revealed that colonies from two coral species, *Porites* sp. and *Pocillopora* sp., that were exposed to high environmental variability generated by internal waves, were mostly immune to acute heat stress treatments. In contrast, their conspecifics from sheltered shores with low environmental variability demonstrated thermal sensitivity under these treatments. Importantly, we showed that stress-hardening through environmental variability can exist under a stimulus comprising low temperature pulses (down to ∼26.5°C and minima of 24°C) that fluctuate at large amplitudes (∼ 3°C ranging between 0.7 - 5.4°C). Secondly, we found that the thermal resistance of corals from exposed reefs persisted throughout the year in presence and in absence of the internal wave stimulus. However, thermal stress resistance levels appeared to fade slightly during the season when the stimulus was absent. Surprisingly, the dynamics of thermal stress resistance were very similar in both coral species, *Pocillopora* sp. and *Porites* sp. despite representing a naturally thermo-sensitive and a robust coral ecotype, respectively (Brown and Phongsuwan 2012; Schmidt et al. 2012; Jain et al. 2023). In the following we discuss these new insights about environmental priming, while considering the regional context of the Thai Andaman Sea, and with regard to designing efficient preconditioning treatments that can enhance coral thermal resistance for coral reef conservation.

### Pulses of cooler temperatures and large amplitudes of variation provide a stress-hardening stimulus for corals in the Thai Andaman Sea

Various organisms, including corals (Rivest et al. 2017; Hackerott et al. 2021)), are known to be more stress-tolerant, when previously exposed to environmental variability (Nicotra et al. 2010; Li et al. 2014; Hilker et al. 2016; Hilker and Schmülling 2019). However, the underlying environmental drivers that induce environmental variability and generate such “stress-hardening regimes” in coral reefs differ between habitat types and reef locations. Since temperature is a major determinant of coral reef distribution and warming poses a threat to corals (Hoegh-Guldberg 1999; Kleypas et al. 1999), it has been commonly used as a proxy to characterize reefs and quantify their environmental variability (Leichter et al. 1996; Oliver and Palumbi 2011a; Wall et al. 2012; Kenkel et al. 2015). In this regard, the most commonly investigated stimuli for stress hardening were elevated and/or fluctuating temperatures that are a feature of shallow reef flats, tidal pool sites or lagoon-type habitats (Palumbi et al. 2014; Camp et al. 2016, 2017). In these locations, corals experience temperature conditions that often exceed the local bleaching thresholds during midday which provide “training periods” for more severe heat wave conditions. It seems intuitive that corals exposed to such conditions, occurring in short term-intervals, “learn” to cope with the environmental stress of elevated temperatures that would usually lead to massive bleaching events.

Notably, the environmental variability regime in our study sites was induced by internal waves and can be so far considered a unique scenario, as it differs from the other locations where environmental priming has been typically investigated. In our study area, reef sites on the western shores of the islands were exposed to the physical forces of internal waves, which create the remarkable difference in environmental conditions between the exposed, western reef sites and the sheltered, eastern sites (Schmidt et al. 2012; Wall et al. 2012, 2015). Internal waves in the Andaman Sea transport cooler waters from the depths to the reefs, significantly increasing the amplitude of temperature variation on western island shores. Other than in reef flat or tidal pool sites where others have investigated environmental priming, internal waves at our study sites provided stimuli of lower rather than higher temperatures. Irrespective of this thermal difference, we report that corals exposed to internal waves were able to cope better with short-term acute heat stress conditions than corals living without the stimulus of internal waves. We propose that the specific amplitude of variability (∼3-5°C) in our west-shore study sites and the lowest temperatures, likely reaching lower thermal threshold (minima of 24°C), might be equally relevant for environmental priming, as established for fluctuating temperatures that temporarily transgress upper thermal thresholds (Oliver and Palumbi 2011a).

Indeed, short pulses of cold water have been shown to induce an acute stress response in corals, but corals could more easily acclimatize to the cooling treatment in comparison to the heat treatment where coral health slowly declined (Roth et al. 2012). In another study, corals performed slightly better under a cooler but “sublethal” temperature compared to the ambient corals, being able to build up more mass and energy reserves (Nielsen et al. 2020). Based on these reports and our new insights, we propose that the effects of low temperature pulses deserve to be further investigated, as they could offer a stress-hardening regime that might emerge as more efficient than the application of high temperature pulses. This could entail accumulation of heat stress when thermal thresholds are exceeded, leading to a negative effect.

To date, findings supporting the effect of environmental priming regimes on coral thermal tolerance are equivocal. While most studies indicate that a “challenging” thermal history or preconditioning regime (of thermal variability or elevated baseline temperature) enhances thermal tolerance of corals (McClanahan et al. 2005; Bellantuono et al. 2012b; Palumbi et al. 2014; Buerger et al. 2015; Schoepf et al. 2015; Kenkel and Matz 2016; DeMerlis et al. 2022; Brown et al. 2023), some report neutral or negative results, including cases where corals ended up less stress resistant compared to the control group (Putnam and Edmunds 2011; Camp et al. 2016; Schoepf et al. 2019; Henley et al. 2022). It has been suggested that such preconditioning treatments must have exerted too much stress on the corals with the consequence of having drained their energy reserves, hence did not contribute to stress-hardening but rather had a contrary effect (Hackerott et al. 2021; Wong et al. 2021). As such, regimes of variability at elevated temperatures can be difficult to implement. We often have a limited understanding of the thermal performance curve of corals, in particular in regard to their species- and location-specific thermal optimum, as well as to their upper critical temperature (Sinclair et al. 2016; Hillebrand et al. 2020). This impedes the determination of an effective environmental regime that can achieve a positive effect of stress-hardening. Considering our knowledge about the effects of cooler temperature on corals and the results in our study, we conclude that cold-stress could be an effective tool to stress-harden corals, as it can successfully trigger metabolic flexibility without the effect of stress accumulation through depletion of energy reserves or taxing coral symbionts.

### Seasonality of the stimulus and the durability of the environmental priming effect

Internal wave activity is seasonal in the shallow reef habitat of the Andaman Sea (Schmidt et al. 2012; Wall et al. 2012). During the first part of the year internal waves expose corals to cold, deep water at regular intervals. At that time, the western shore is usually hit by the waves and thus the stimulus is at its peak, creating the largest environmental differences between the western and the sheltered, eastern shore. Later in the year, internal wave impact dwindles and consequently the environmental conditions on both island shores, east and west, become very similar. We took advantage of this seasonality in the Andaman Sea region to explore the effects of the presence and absence of a variable stimulus. To date, the persistence of a stress-hardening effect in corals has hardly been considered in great detail and remains to be investigated (Klepac and Barshis 2022). Our results showed that the positive effect on thermal resistance had prevailed even in the absence of the stimulus. However, we observed a slight, but measurable decline of stress resistance during the season of stimulus absence. This speaks for the case that the effect of stress-hardening is lasting, but could slowly fade in the complete absence of the priming stimulus. Our observation aligns with the finding that bleaching thresholds of corals decreased seasonally, e.g., during the cooler winter season, when thermal challenges of the summer time were absent (Berkelmans and Willis 1999). Similarly, this has been the case for one Caribbean coral species (Scheufen et al. 2017), however, the same study found that other coral species did not follow this seasonality. The latter agrees with several other cases which have shown that corals maintained their stress resistance levels after transplantation from a high variability reef to an aquarium or site with more stable conditions (Morikawa and Palumbi 2019; Schoepf et al. 2019; Marhoefer et al. 2021). Also, the effect of various thermal preconditioning treatments (variable and stable) had a measurable effect on coral thermal tolerance four months later (Drury et al. 2022). Notably, Morikawa and Palumbi (2020) have observed the permanence of stress resistance across coral taxa in their coral nursery hosting resilient corals from high variability sites for two consecutive years.

### Disentangling covariates of variable environmental priming regimes

Reef sites that are exposed to internal waves have an important advantage over the typically investigated intertidal and lagoon-type coral habitats. They represent a reef habitat with all typical physicochemical features of an ocean-facing reef slope and can be compared to similar reefs that are sheltered from the impact of internal waves. This provides a setting where the effects of reef site-specific characteristics are accounted for and effects induced by the variability regime can be studied in isolation. In contrast, tidal pools and lagoons are habitats that are fundamentally different from the typical coral reef. Extreme light intensities and elevated salinities are characteristic for these shallow sites (Yates et al. 2014). Yet, these sites are often compared to control sites located in a proper reef slope habitat, which does not allow to disentangle the sole effect of the variability experienced in these sites. Still, it is important to consider that corals in our study were not solely challenged by temperature fluctuations caused by the internal waves. The deep-water brought into the reefs by internal waves is also typically rich in inorganic nutrients and particulate matter. Both can be either beneficial or challenging for corals and microalgal symbionts (Risk 2014). On one hand, the increase in nutrient sources could be valuable for corals and contribute to their resilience (Ferrier-Pagès et al. 2000; Meunier et al. 2022). On the other hand, particle loads may reduce light penetration and reduce photosynthetic output of microalgal symbionts (Anthony et al. 2007). In addition, sediment particles typically threaten to smother corals (Tuttle et al. 2020) that will need to spend energy on mucus production to free their tissues from these sediments. Depending on the amount of nutrients introduced by internal waves and the requirements of the corals, increases in inorganic nutrients and particulates can lead to a nutrient imbalance that can threaten the intricate balance between host and algal symbiont (Wiedenmann et al. 2012; Rädecker et al. 2015; Morris et al. 2019). Similarly, a slightly lower pH and oxygen-depleted seawater carried by internal waves (Schmidt et al. 2012; Wall et al. 2012) may pose a stressor to corals and challenge their performance (Chan and Connolly 2013; Alderdice et al. 2021). To better understand how co-variation of these variables influences coral physiology and stress-hardening in the Andaman Sea, holistic surveys will be needed that assess and consider a diversity of physico-chemical variables. However, this is not only important when studying internal wave sites. Co-fluctuating variables exist in all types of high variability reef sites, including tidal pools, reef flats, and lagoons in proximity to seagrass or mangroves (Ruiz-Jones and Palumbi 2015; Camp et al. 2016). To explain some of the ambiguous findings of coral stress-hardening studies, future research will need to explore the effects of co-varying variables, as they may play a role in modulating the effects of stress-hardening.

### Considerations for the design of efficient stress-hardening regimes

Studies of stress-hardening in corals through thermal variability regimes have sparked the idea of instrumentalizing this phenomenon to improve coral thermal stress resistance during climate change. It is anticipated that simulation of a stress-hardening stimulus can be used to enhance thermal tolerance of corals for the purpose of conservation and restoration of coral reefs (Middlebrook et al. 2008; Bellantuono et al. 2012b). The phenomenon, however, is still poorly understood and some findings remain equivocal. While many studies have reported positive effects of a variable environment on the stress tolerance in corals (Doering et al. 2021; Oliver and Palumbi 2011b; Buerger et al. 2015; Wong et al. 2021; DeMerlis et al. 2022; Brown et al. 2023), a few have not reported any improvements or rather observed declines in stress tolerance. Negative reports are likely due to stress-buildup during the preconditioning process (Hackerott et al. 2021), which can occur when a variability regime becomes too challenging (Putnam and Edmunds 2011; Camp et al. 2016; Schoepf et al. 2019; Klepac and Barshis 2020; Henley et al. 2022). Also, dynamic interaction of all covariates present in the respective study sites can act as confounding factors and influence the outcomes of preconditioning (as laid out in the chapter above), but most importantly, the “priming dosage” will be decisive for the success of the method. Fine-scale differences in the amplitude and frequency of variation employed (Klepac and Barshis 2022; Brown et al. 2023), the average temperature in the preconditioning regime, as well as the duration of the exposure (Bellantuono et al. 2012a; Hackerott et al. 2021; Martell 2023) deserve careful consideration. In some studies that have failed to observe a positive effect, environmental variability in the reef sites or treatments might have been too small in comparison to the ambient regime in order to elicit a measurable effect on corals. For instance, the variability ranges of the study sites in (Camp et al. 2016), only differed by ∼1 - 2°C, which might be too small of a difference to pinpoint any effect (Rivest et al. 2017). Only a few efforts so far have set out to systematically identify optimal priming regimes. Early surveys and experiments have found that heat tolerance was correlated with the magnitude of variability, as corals from the tidal pool with the highest variability appeared to be most resistant to heat (Palumbi et al. 2014). Several recent study designs have allowed us to gain insights at a higher resolution and have found that an intermediate variability regime might likely be the most effective for stress-hardening of corals. For instance, corals living in sites of intermediate variability on Heron Island in the Great Barrier Reef (Brown et al. 2023) or in the moderately variable pools of the well-known study sites in American Samoa (Klepac and Barshis 2022), have outperformed conspecifics that had experienced lower or higher variability. Most recent findings suggest that exposure to thermal variability at a rather low average mean temperature, or involving cooling rather than heat pulses, could be more efficient than variability at a higher average temperature, as it has led to better stress-hardening results in corals (Drury et al. 2022). Future studies will be needed to further refine our understanding of how environmental priming regimes work, which will lead to the design of efficient preconditioning protocols.

On a last note, it still remains to be elucidated whether and at which cost(s) stress-hardened corals acclimate to perform well under challenging environmental regimes. Trade-offs will be an important aspect of future investigations. Conservation and restoration efforts that aim to apply preconditioning strategies to stress-harden corals will need to evaluate whether the gain in thermal resistance is related to any critical trade-offs. At our study sites in the Andaman Sea, resistance of west-shore corals might be coupled with a lower reef framework building capacity that was reported from these sites earlier (Schmidt et al. 2012; Wall et al. 2012). This calls for detailed investigations into the calcification capacity of these resistant corals. Recent study focussing on trade-offs (Wong et al. 2021) have found that corals from high variability sites or long-term high-temperature treatments, had either a lower metabolic capacity, lower growth rates, or lower reproductive potential compared to the control groups from stable or ambient habitats or treatments. Overall, efforts aimed at increasing thermal tolerance of corals will need a holistic approach to the subject. For the development of new interventions, it will be essential to carefully assess cost-benefits and evaluate each new method and its potential ecological consequences.

## Conclusion

We showed that two coral species that occupy different ecological niches were receptive to the same environmental priming of cooling pulses, which improved their thermal stress resistance to acute short-term heat stress. A cold-water priming pulse can induce stress-hardening effectively. It might be a safer option compared to the implementation of high temperature peaks in variability regimes used as preconditioning treatments as heat-stress accumulation is avoided. Our study also showed that a temporary priming exposure can induce a stress-hardening effect, which, however, is likely to fade in longer absence of the stimulus, suggesting that a reapplication of a preconditioning treatment will be necessary. Most importantly, the ideal dosage and length of thermal variability exposure in a preconditioning treatment will need to be determined. Eventually, the enhancement of stress resistance traits is likely to come at the cost of other traits. Therefore, research into the trade-offs that accompany thermal resistance gain in corals will be crucial in order to understand the capacity and limitations of corals to resist future thermal stress.

## Supporting information

Wall_Roik_Et_Al_stresshardening_corals

## Acknowledgments

We acknowledge the team of the research unit Marine Ecology at Phuket Marine Biological Center for field logistics and support during the set up of coral experimental facilities. We thank M. Heckwolf, K. Bimson, L. Niewendieck, M. Suwareh, and V. Conrad for field assistance. We thank F. Wendt for advice regarding aquarium equipment. This research was funded by the DFG (German National Science Foundation) excellence initiative “Future Ocean” (# CP1782) awarded to AR. A.R. further supported by funding of the Helmholtz Institute for Functional Marine Biodiversity at the University of Oldenburg, Niedersachsen, Germany. HIFMB is a collaboration between the Alfred-Wegener-Institute, Helmholtz-Center for Polar and Marine Research, and the Carl-von-Ossietzky University Oldenburg, initially funded by the Ministry for Science and Culture of Lower Saxony and the Volkswagen Foundation through the “Niedersächsisches Vorab” grant program (grant number ZN3285).

## Permissions

Research in Thailand was conducted under the permit of the National Research Council of Thailand (NRCT # 0002/632), and corals were collected under the collection permission of CITES (export # AC.0510.6/0017 and # AC.0510.6/022; import # DE E-04829/18 and # DE E-01510/19).

## Authors’ contributions

AR conceived the study. AR and TD designed the experiment. TD, AR, MW conducted coral experiments. AR, MW, LP, and TR performed coral collection. TD, NP, MW, AR performed the data analysis. AR generated data visualization. AR, MW, TD wrote and edited the manuscript. Field facilities and logistics were provided by LP, TR and the PMBC team. The authors read and approved the final manuscript.

## Declarations

The authors declare that they have no competing interests.

