## Supplementary material for "Natural thermal stress-hardening of corals through cold temperature pulses in the Thai Andaman Sea": Wall_Roik_Et_Al_stresshardening_corals

### Supplementary Materials

#### Text S1

The coral maintenance facility was located at Phuket Marine Biological Center (Cape Panwa, Phuket, Thailand) within an hour by speed boat from the most distant sampling site. Collected coral fragments (*Pocillopora*: length ~ 5 cm; *Porites*: ø ~ 6 cm) were maintained in two large 500 L flow-through tanks with a flow rate of 2.8 ± 1.31 L/min and the average ambient *in situ* temperature of the season for 2-12 days before used in the experiments. A 500 L source tank constantly supplied 5 µm-filtered seawater from the reef adjacent to the facilities and temperature was held at constant 29.43 ± 0.32 °C using a chiller, a heater, and a temperature-control device (Aqua Medic Titan 1500 Chiller, Germany; Titanium Heater 100 W, Schego, Germany; Temperature Switch TS 125, HTRONIC, Germany). During experiments, i.e., heat tolerance assessment, the large flow-through tanks served as temperature-stabilizing baths for four 40 L experimental tanks, two per water bath. Experimental tanks were supplied through daily manual water change (twice daily 50%) from the source tank. Each experimental tank was equipped with a temperature-control device, one heater, air supply, a small current pump, and a temperature logger (Titanium Heater 100 W, Schego, Germany; Temperature Switch TS 125, HTRONIC, Germany; HOBO Pendant Temperature/Light 8K Data Logger, Onset, USA; Koralia nano 900 L/h, Hydor, Italy). Each of the four experimental tanks as well as the flow-through tanks were equipped with LED lights (135 W, Hydra Fiftytwo HD LED, Aqua Illumination, USA) that mimicked the average light conditions of the sampling sites. Tanks were monitored regularly by measuring a suite of environmental parameters (temperature, oxygen, light intensity, and salinity). Briefly, temperature was measured continuously with loggers (HOBO Pendant® Temperature/Light 64K Data Logger, Onset, USA). Other parameters were monitored at regular time intervals, i.e., photosynthetically active radiation (PAR) measured by a quantum meter (MQ-210 Underwater Quantum Meter, Apogee Instruments, USA), dissolved oxygen and salinity measured by a hand-held multimeter (Multi3430, FDO®925, and TetraCon®925, WTW, Germany). Physico-chemical parameters of tank conditions are provided in Tables S2.

**Table S1 Replicate numbers of coral colonies tested per reef sites and seasons.**

| Coral species | Site of origin | Season of stimulusS presence | Season of stimulusS absence |
|---|---|---|---|
| *Porites* sp. | West \| High variability site | N=12 | N=12 |
| | East \| Low variability site | N=13 | N=14 |
| *Pocillopora* sp. | West \| High variability site | N=10 | N=18 |
| | East \| Low variability site | N=9 | N=13 |

*stimulus = environmental priming stimulus of high thermal variability generated by internal waves

**Table S2 Summary of tank conditions during the heat stress assays.** Temperature, oxygen, light intensity and salinity data (mean ± SD) are presented for the duration of the assay for both coral species. Temperature is specifically summarized for the temperature-peak period for each treatment (i.e., '29 °C' and '34 °C').

| Coral species | Heat stress assay treatments | Temperature (°C) | Peak temperature (°C) | Dissolved oxygen (mg/L) | Light intensity (µmol $m^{-2}$ $s^{-2}$) | Salinity (PSU) |
|---|---|---|---|---|---|---|
| *Pocillopora* sp. | *'29 °C'* | 29.82 ± 0.35 | 30.08 ± 0.19 | 7.61 ± 0.02 | 74 ± 3 | 32.7 ± 0.1 |
| | *'34 °C'* | 30.99 ± 2.03 | 33.89 ± 0.56 | 7.63 ± 0.01 | 75 ± 5 | 32.4 ± 0.1 |
| *Porites* sp. | *'29 °C'* | 29.14 ± 0.18 | 29.35 ± 0.18 | 8.20 ± 0.27 | 83 ± 8 | NA |
| | *'34 °C'* | 30.46 ± 2.01 | 34.27 ± 0.16 | 8.16 ± 0.29 | 84 ± 9 | NA |

**Table S3 Effect sizes of coral stress responses.** Effect sizes are represented by the mean differences (response under ambient temperature minus response under heat treatment) calculated through bootstrap estimation (*dabestR* R-package). The 95% confidence intervals (95CI) are indicated in square brackets. 5000 bootstrap resamples were used and CIs are bias-corrected and accelerated. Negative values indicate the decline of holobiont tissue coloration (i.e., bleaching score) and symbiont photosynthetic efficiency (i.e., Effective Quantum Yield).

| | | Δ Bleaching Score | | Δ Effective Quantum Yield | |
|---|---|---|---|---|---|
| **Site of origin** | **Season of** | ***Pocillopora* sp.** | ***Porites* sp.** | ***Pocillopora* sp.** | ***Porites* sp.** |
| West \| High variability site | Stimulus presence | -0.5 [95CI -1.06; -0.0556] | -0.667 [95CI -1.17; -0.167] | 0.001 [95CI -0.016; 0.022] | -0.00656 [95CI -0.041; 0.020] |
| West \| High variability site | Stimulus absence | -0.806 [95CI -1.25; -0.444] | -0.583 [95CI -0.958; -0.292] | -0.0362 [95CI -0.060; -0.022] | -0.0193 [95CI -0.027; -0.012] |
| East \| Low variability site | Stimulus presence | -1.05 [95CI -1.75; -0.3] | -1.97 [95CI -2.63; -1.4] | -0.0246 [95CI -0.045; -0.009] | -0.0289 [95CI -0.078; 0.007] |
| East \| Low variability site | Stimulus absence | -1.77 [95CI -2.23; -1.38] | -1.14 [95CI -1.5; -0.75] | -0.041 [95CI -0.060; -0.024] | -0.0339 [95CI -0.048; -0.022] |

**Table S4 Generalized Linear Mixed-Effects Models.** Models were built based on the formula *data~site*treatment+(1*site|colony)* employing Gaussian or Gamma distribution. Eight models were generated, one per response variable (bleaching score and photosynthetic performance) for each coral species per season. data = full data set; data_rmout2 = data with outliers removed.

| **Response variable** | **Season** | **Cora species** | **Data** | **shapiro *p*** | **AIC** | **BIC** | **logLik** | **deviance** | **df.resid** | **Model** | **Distribution** |
|---|---|---|---|---|---|---|---|---|---|---|---|
| Bleaching Score | Stimulus presence | Pocillopora | "data" | 0.096 | **94.4** | 104.2 | -41.2 | 82.4 | 32 | glmerMod | Family: gaussian ( identity ) |
| | | Porites | "data" | 0.021 | 141.8 | 153.7 | -64.9 | 129.8 | 48 | glmerMod | Family: Gamma ( log ) |
| | Stimulus absence | Pocillopora | "data" | 0.000 | **128** | 140.8 | -58 | 116 | 56 | glmerMod | Family: Gamma ( log ) |
| | | Porites | "data" | 0.001 | **80.4** | 92.1 | -34.2 | 68.4 | 46 | glmerMod | Family: Gamma ( log ) |
| Photosynthetic Efficiency | Stimulus presence | Pocillopora | "data_rmout2" | 0.004 | -182.1 | -172.6 | 97.1 | -194.1 | 30 | glmerMod | Family: Gamma ( log ) |
| | | Porites | "data" | 0.000 | -155.1 | -143.2 | 83.5 | -167.1 | 48 | glmerMod | Family: Gamma ( log ) |
| | Stimulus absence | Pocillopora | "data_rmout2" | 0.000 | -300 | -287.4 | 156 | -312 | 54 | glmerMod | Family: Gamma ( log ) |
| | | Porites | "data" | 0.000 | -282.8 | -271.1 | 147.4 | -294.8 | 46 | glmerMod | Family: Gamma ( log ) |

**Table S5 Fixed effects reported from generalized Linear Mixed-Effects Models for the bleaching score response variable.** Statistics and *p*-values are reported for each coral species. Site = reef site of origin (East, West); treatment = treatment group of heat stress assay (ambient temperature control group, heat stress group 34 °C)

| ***Pocillopora* \| Season of stimulus presence** | **Estimate** | **Std. Error** | **t value** | **Pr(>\|z\|)** | **Significance level** |
|---|---|---|---|---|---|
| ## (Intercept) | 10.20152 | 0.22569 | 45.202 | < 2e-16 | *** |
| ## site | 0.04221 | 0.30743 | 0.137 | 0.890805 | |
| ## treatment | -1.05001 | 0.29206 | -3.595 | 0.000324 | *** |
| ## site:treatment | 0.52232 | 0.42828 | 1.22 | 0.222621 | |
| ***Porites* \| Season of stimulus presence** | **Estimate** | **Std. Error** | **t value** | **Pr(>\|z\|)** | **Significance level** |
| (Intercept) | 2.35095 | 0.02267 | 103.72 | < 2e-16 | *** |
| site | -0.01902 | 0.0311 | -0.612 | 0.54079 | |
| treatment | -0.20899 | 0.02901 | -7.204 | 5.85E-13 | *** |

| | | | | | |
|---|---|---|---|---|---|
| site:treatment | 0.1419 | 0.0435 | 3.262 | 0.00111 | ** |
| ***Pocillopora* \| Season of stimulus absence** | **Estimate** | **Std. Error** | **t value** | **Pr(>\|z\|)** | **Significance level** |
| (Intercept) | 2.34647 | 0.01831 | 128.173 | < 2e-16 | *** |
| site | -0.01468 | 0.02334 | -0.629 | 0.52942 | |
| treatment | -0.18671 | 0.02375 | -7.862 | 3.78E-15 | *** |
| site:treatment | 0.10486 | 0.03114 | 3.367 | 0.00076 | *** |
| ***Porites* \| Season of stimulus absence** | **Estimate** | **Std. Error** | **t value** | **Pr(>\|z\|)** | **Significance level** |
| (Intercept) | 2.27299 | 0.0147 | 154.663 | < 2e-16 | *** |
| site | 0.02266 | 0.01825 | 1.242 | 0.2144 | |
| treatment | -0.12531 | 0.01736 | -7.218 | 5.28E-13 | *** |
| site:treatment | 0.06478 | 0.02555 | 2.535 | 0.0112 | * |

**Table S6 Fixed effects reported from generalized Linear Mixed-Effects Models for the photosynthetic efficiency variable.** Statistics and *p*-values are reported for each coral species. Site = reef site of origin (East, West); treatment = treatment group of heat stress assay (ambient temperature control group, heat stress group 34 °C)

| ***Pocillopora* \| Season of stimulus presence** | **Estimate** | **Std. Error** | **t value** | **Pr(>\|z\|)** | **Significance level** |
|---|---|---|---|---|---|
| (Intercept) | 0.6925552 | 0.0048485 | 142.839 | < 2e-16 | *** |
| site | 0.0001751 | 0.0031736 | 0.055 | 0.95599 | |
| treatment | -0.0123004 | 0.003027 | -4.064 | 4.83E-05 | *** |
| site:treatment | 0.0112807 | 0.0042945 | 2.627 | 0.00862 | ** |
| ***Porites* \| Season of stimulus presence** | **Estimate** | **Std. Error** | **t value** | **Pr(>\|z\|)** | **Significance level** |
| (Intercept) | 0.678404 | 0.007378 | 91.949 | <2e-16 | *** |
| site | 0.011163 | 0.009615 | 1.161 | 0.2456 | |
| treatment | -0.014834 | 0.008922 | -1.663 | 0.0964 | . |
| site:treatment | 0.011486 | 0.013382 | 0.858 | 0.3907 | |

| *Pocillopora* \| Season of stimulus absence | Estimate | Std. Error | t value | Pr(>\|z\|) | Significance level |
|---|---|---|---|---|---|
| (Intercept) | 0.6936882 | 0.0027356 | 253.582 | < 2e-16 | *** |
| site | -0.0006219 | 0.003258 | -0.191 | 0.849 | |
| treatment | -0.0206975 | 0.0033106 | -6.252 | 4.05E-10 | *** |
| site:treatment | 0.0064631 | 0.0043289 | 1.493 | 0.135 | |
| ***Porites* \| Season of stimulus absence** | **Estimate** | **Std. Error** | **t value** | **Pr(>\|z\|)** | **Significance level** |
| (Intercept) | 0.689365 | 0.002219 | 310.641 | < 2e-16 | *** |
| site | 0.002844 | 0.002583 | 1.101 | 0.2709 | |
| treatment | -0.017184 | 0.002453 | -7.005 | 2.48E-12 | *** |
| site:treatment | 0.007466 | 0.003611 | 2.068 | 0.0387 | * |

**Table S7 Post Hoc comparisons for bleaching score data.** Statistics and *p*-values are reported for each coral species per season. Reef sites of origin: RE = East shore reef, RW = West shore reef; Groups of heat stress assays: A = ambient temperature control group, H = heat stress group 34 °C.

| contrast | estimate | SE | df | *z*-ratio | *p*-value |
|---|---|---|---|---|---|
| ***Pocillopora* \| Season of stimulus presence** | | | | | |
| RE A - RW A | -0.0422 | 0.307 | Inf | -0.137 | 0.9991 |
| **RE A - RE H** | **1.05** | **0.292** | **Inf** | **3.595** | **0.0018** |
| RE A - RW H | 0.4855 | 0.306 | Inf | 1.584 | 0.3875 |
| **RW A - RE H** | **1.0922** | **0.307** | **Inf** | **3.553** | **0.0022** |
| RW A - RW H | 0.5277 | 0.313 | Inf | 1.685 | 0.3318 |
| RE H - RW H | -0.5645 | 0.306 | Inf | -1.842 | 0.2534 |
| ***Porites* \| Season of stimulus presence** | | | | | |
| contrast | estimate | SE | df | z.ratio | p.value |
| RE A - RW A | 0.019 | 0.0311 | Inf | 0.612 | 0.9284 |
| **RE A - RE H** | **0.209** | **0.029** | **Inf** | **7.204** | **<.0001** |

| | | | | | |
|---|---|---|---|---|---|
| **RE A - RW H** | **0.0861** | **0.0311** | **Inf** | **2.768** | **0.0289** |
| **RW A - RE H** | **0.19** | **0.0311** | **Inf** | **6.111** | **<.0001** |
| RW A - RW H | 0.0671 | 0.0324 | Inf | 2.07 | 0.1631 |
| **RE H - RW H** | **-0.1229** | **0.0311** | **Inf** | **-3.952** | **0.0005** |
| ***Pocillopora* \| Season of stimulus absence** | | | | | |
| RE A - RW A | 0.0147 | 0.0233 | Inf | 0.629 | 0.9228 |
| **RE A - RE H** | **0.1867** | **0.0237** | **Inf** | **7.862** | **<.0001** |
| **RE A - RW H** | **0.0965** | **0.0235** | **Inf** | **4.109** | **0.0002** |
| **RW A - RE H** | **0.172** | **0.0232** | **Inf** | **7.424** | **<.0001** |
| **RW A - RW H** | **0.0818** | **0.0202** | **Inf** | **4.057** | **0.0003** |
| **RE H - RW H** | **-0.0902** | **0.0233** | **Inf** | **-3.87** | **0.0006** |
| ***Porites* \| Season of stimulus absence** | | | | | |

| | | | | | |
|---|---|---|---|---|---|
| RE A - RW A | -0.0227 | 0.0183 | Inf | -1.242 | 0.6003 |
| **RE A - RE H** | **0.1253** | **0.0174** | **Inf** | **7.218** | **<.0001** |
| RE A - RW H | 0.0379 | 0.0183 | Inf | 2.074 | 0.1615 |
| **RW A - RE H** | **0.148** | **0.0182** | **Inf** | **8.109** | **<.0001** |
| **RW A - RW H** | **0.0605** | **0.0187** | **Inf** | **3.228** | **0.0068** |
| **RE H - RW H** | **-0.0874** | **0.0183** | **Inf** | **-4.79** | **<.0001** |

**Table S8 Post Hoc comparisons for photosynthetic efficiency data.** Statistics and *p*-values are reported for each coral species per season. Reef sites of origin: RE = East shore reef, RW = West shore reef; Groups of heat stress assays: A = ambient temperature control group, H = heat stress group 34 °C.

| contrast | estimate | SE | df | z.ratio | p.value |
|---|---|---|---|---|---|
| ***Pocillopora* \| Season of stimulus presence** | | | | | |
| RE A - RW A | -0.00018 | 0.00317 | Inf | -0.055 | 0.9999 |
| **RE A - RE H** | **0.0123** | **0.00303** | **Inf** | **4.064** | **0.0003** |

| | | | | | |
|---|---|---|---|---|---|
| RE A - RW H | 0.000845 | 0.00314 | Inf | 0.269 | 0.9932 |
| **RW A - RE H** | **0.012475** | **0.00333** | **Inf** | **3.743** | **0.001** |
| RW A - RW H | 0.00102 | 0.00307 | Inf | 0.333 | 0.9873 |
| **RE H - RW H** | **-0.01146** | **0.00328** | **Inf** | **-3.49** | **0.0027** |
| ***Porites* \| Season of stimulus presence** | | | | | |
| RE A - RW A | -0.01116 | 0.00961 | Inf | -1.161 | 0.6515 |
| RE A - RE H | 0.01483 | 0.00892 | Inf | 1.663 | 0.3436 |
| RE A - RW H | -0.00781 | 0.00961 | Inf | -0.813 | 0.8485 |
| **RW A - RE H** | **0.026** | **0.00961** | **Inf** | **2.705** | **0.0345** |
| RW A - RW H | 0.00335 | 0.00997 | Inf | 0.336 | 0.987 |
| RE H - RW H | -0.02265 | 0.00961 | Inf | -2.356 | 0.0857 |
| ***Pocillopora* \| Season of stimulus absence** | | | | | |

| | | | | | |
|---|---|---|---|---|---|
| RE A - RW A | 0.000622 | 0.00326 | Inf | 0.191 | 0.9975 |
| **RE A - RE H** | **0.020697** | **0.00331** | **Inf** | **6.252** | **<.0001** |
| **RE A - RW H** | **0.014856** | **0.00332** | **Inf** | **4.473** | **<.0001** |
| **RW A - RE H** | **0.020076** | **0.00332** | **Inf** | **6.041** | **<.0001** |
| **RW A - RW H** | **0.014234** | **0.00279** | **Inf** | **5.104** | **<.0001** |
| RE H - RW H | -0.00584 | 0.00339 | Inf | -1.725 | 0.3104 |
| ***Porites* \| Season of stimulus absence** | | | | | |
| RE A - RW A | -0.00284 | 0.00258 | Inf | -1.101 | 0.6889 |
| **RE A - RE H** | **0.01718** | **0.00245** | **Inf** | **7.005** | **<.0001** |
| **RE A - RW H** | **0.00687** | **0.00258** | **Inf** | **2.661** | **0.039** |
| **RW A - RE H** | **0.02003** | **0.00258** | **Inf** | **7.752** | **<.0001** |
| **RW A - RW H** | **0.00972** | **0.00265** | **Inf** | **3.667** | **0.0014** |

| | | | | | |
|---|---|---|---|---|---|
| **RE H - RW H** | **-0.01031** | **0.00258** | **Inf** | **-3.991** | **0.0004** |
| **RE H - RW H** | **-0.01031** | **0.00258** | **Inf** | **-3.991** | **0.0004** |
